# Therapeutic knockdown of MLKL reduces diet-induced obesity and improves insulin signalling in mature adipocytes

**DOI:** 10.64898/2026.04.17.719119

**Authors:** Masaaki Sato, Xichun Li, Han Xu, Abdulaziz MA Alammar, Shevin C Fernando, Marziyeh Anari, Komal Patel, Kritima Dhakal, Sidney Niogret, Yizhuo Wang, Taaseen Rahman, Yung-Chih Chen, Stephen J Nicholls, Brian G Drew, James M Murphy, Denuja Karunakaran

## Abstract

Obesity affects one in three adults and is complicated by adipose inflammation, lipotoxicity and cell death. We previously identified RIPK1 as a genetic determinant of human obesity risk and adipose inflammation. Because RIPK1 is the apical kinase in the necroptosis pathway upstream of RIPK3 and the executioner protein MLKL, and emerging evidence links MLKL to lipid metabolism, MLKL has surfaced as a potential metabolic regulator. However, conflicting findings in *Mlkl* knockout mice fed a high fat diet have left its therapeutic relevance unresolved. MLKL has not been previously targeted through therapeutic knockdown *in vivo* in the context of diet-induced obesity. Here, we evaluated two independent MLKL antisense oligonucleotides (ASOs) in high fat diet (HFD)-fed C57BL/6J mice. In a 24-week progression model, MLKL ASO markedly reduced body weight, fat mass and hepatic steatosis compared with controls, while preserving lean mass. MLKL knockdown also lowered the respiratory exchange ratio, indicating a shift toward increased fat oxidation. In the intervention model, once obesity was established after 12 weeks of HFD feeding, both MLKL ASOs, and similarly, two independent RIPK1 ASOs, reversed weight gain and improved systemic glucose control. *In vitro*, MLKL-CRISPR/Cas9 knockout blocked 3T3-L1 adipogenesis, indicating a requirement for MLKL during adipocyte differentiation. However, in mature adipocytes, MLKL siRNA reduced palmitic acid-induced lipid accumulation, increased isoprenaline-stimulated lipolysis, and prevented TNF-mediated suppression of insulin-mediated AKT signalling and glucose uptake. Collectively, these findings demonstrate that partial MLKL suppression reprograms whole-body energy metabolism, enhances insulin sensitivity and limits diet-induced adiposity. MLKL, therefore, represents a promising and mechanistically novel therapeutic target for obesity and insulin resistance.

## INTRODUCTION

Obesity and its associated metabolic complications, including insulin resistance, type 2 diabetes, and metabolic dysfunction-associated steatotic liver disease (MASLD), remain major global health challenges. While excess caloric intake and sedentary behaviour are central drivers, the progression from obesity to metabolic disease is critically shaped by expansion of subcutaneous and visceral adipose depots and adipocyte dysfunction, including impaired lipid metabolism and chronic low-grade inflammation [1, 2]. Current pharmacological approaches, including incretin-based therapies, primarily induce weight loss through appetite suppression, with improvements in insulin sensitivity occurring largely secondary to reduced caloric intake [3]. In parallel with appetite-based interventions, there is continued interest in therapeutic strategies that directly modulate adipocyte-intrinsic pathways regulating lipid metabolism and insulin action.

Necroptosis is a regulated form of inflammatory cell death mediated by receptor-interacting protein kinase-1 (RIPK1), RIPK3, and the executioner protein mixed lineage kinase domain-like protein (MLKL) [4-11]. Although necroptosis has been extensively studied in the context of tissue injury and inflammation [12-14], increasing evidence suggests that components of this pathway exert broader, context-dependent roles beyond lytic cell death. In metabolic tissues, RIPK1 and MLKL have been implicated in cardiometabolic disease, hepatic steatosis, diet-induced obesity or diabetes [15-21], though their precise functions remain incompletely defined and, in some cases, controversial [22]. Emerging data indicate that MLKL may influence metabolic homeostasis through necroptosis-independent mechanisms [15, 18, 19]. Several studies report that MLKL deficiency protects against diet-induced obesity and hepatic steatosis [17-19, 23], whereas others observe minimal metabolic effects [22], suggesting strong dependence on tissue context, genetic background, and degree of inhibition. In adipose tissue, MLKL expression increases with obesity, and MLKL has been shown to regulate white adipocyte differentiation while sparing beige adipogenesis [23, 24], pointing to a role in adipocyte lipid handling rather than cell death, influencing systemic metabolic health. Excessive triglyceride accumulation within adipocytes promotes lipotoxic stress, inflammation, and insulin resistance, whereas increased lipid turnover and fatty-acid oxidation are associated with improved metabolic flexibility and glucose homeostasis [18, 23, 24]. Further, insulin signalling through the PI3K–AKT pathway coordinates glucose uptake and suppresses lipolysis, and disruption of this pathway by inflammatory cytokines such as tumour necrosis factor (TNF) contributes to adipocyte insulin resistance [25, 26]. However, whether therapeutic suppression of MLKL can reverse established obesity and insulin resistance and whether such effects reflect adipocyte-intrinsic mechanisms remain to be elucidated.

We previously showed that therapeutic RIPK1 inhibition using antisense oligonucleotides (ASOs) reduces NF-κB–driven inflammation and disease burden in mouse models of atherosclerosis and diet-induced obesity [15, 16]. RIPK1 is a key regulator of NF-κB signalling, cell survival, and programmed cell death pathways, including apoptosis and necroptosis [27]. ASOs provide a clinically validated strategy for selective gene silencing in metabolic tissues [28], with several approved therapies (e.g. Nusinersen for muscular atrophy [29]) and additional agents progressing through phase II/III trials to reduce lipids or cardiometabolic complications (e.g. Olezarsen, a GalNAc-conjugated ASO targeting apolipoprotein C-III [30]). We and others have demonstrated efficient delivery of 2′-MOE (2′-O-methoxyethyl) and 2′F-MOE (2′-Fluoro-2′-O-methoxyethyl) ASOs to liver and adipose tissue, and previously showed that RIPK1 knockdown improves insulin sensitivity, reduces adiposity, and remodels adipose immune composition in diet-induced obesity [5].

Given emerging evidence that MLKL regulates lipid metabolism and adipocyte biology beyond its canonical role in necroptosis, we hypothesised that partial MLKL suppression could confer metabolic benefits without inducing overt cell death—an approach not previously tested. This premise is supported by observations that aged MLKL-deficient mice do not exhibit deleterious phenotypes, suggesting that chronic MLKL inhibition is likely to be well tolerated [31]. We therefore investigated whether long-term ASO-mediated MLKL knockdown could reduce adiposity, enhance metabolic flexibility, and improve glucose homeostasis in diet-induced obesity, alongside evaluating the translational potential of targeting RIPK1 or MLKL to treat obesity and insulin resistance.

Herein we demonstrate that therapeutic MLKL silencing in high-fat diet (HFD)-fed mice reduced adiposity, improved hepatic steatosis, and shifted whole-body metabolism toward greater fat oxidation while preserving lean mass. In established obesity, inhibition of either RIPK1 or MLKL reversed weight gain and enhanced glucose–insulin homeostasis. Mechanistic studies showed that MLKL is required for early adipogenesis, whereas its suppression in mature adipocytes reduces lipid accumulation, increases lipolysis, and maintains insulin signalling. Taken together, these findings identify RIPK1 and MLKL as promising therapeutic targets for improving metabolic health in obesity and insulin resistance.

## MATERIALS AND METHODS

### Reagents

DMEM high glucose, GlutaMAX supplement, pyruvate (#10569044), Penicillin-Streptomycin solution (P4458), New bovine calf serum (#16010159), Lipofectamine 3000 (#L3000008), Opti-MEM (#31985062), 2-NBDG (#N13195) and BODIPY 493/503 (#D3821) were purchased from Thermo Fisher Scientific (Australia). Control or MLKL ON-TARGET plus siRNAs ((L-061420-00-0010 or L-01420-00-0005) were obtained from Millennium Science (Australia). Antibodies against phospho-AKT (Ser473; #9271), total AKT (#9272), Vinculin (#13901T), and horseradish peroxidase (HRP)–conjugated anti-rabbit (#7074) and anti-rat (#7077) secondary antibodies were from Cell Signalling Technology (MA, USA). Antibody against MLKL (clone 3H1; #MABC604), fatty-acid free bovine serum albumin (BSA; #126609), palmitic acid (#458440), insulin solution (#I9278) and glycerol assay kit (#MAK117) were obtained from Merck Australia. Clarity Max Enhanced chemiluminescence (ECL; #1705061) were purchased from Bio-Rad Australia and Abcam (USA) respectively. Hoechst 33342 staining solution (#HB0787) was sourced from Genesearch Australia, and recombinant carrier-free mouse TNF (#575206) was obtained from BioLegend (USA).

### Animals

All mouse studies were approved by The University of Queensland Animal Ethics Committee, Office of Research Ethics (approval number IMB/338/19), and conducted in accordance with the Queensland Government Animal Research Act 2001, associated Animal Care and Protection Regulations (2002 and 2008), and the Australian Code for the Care and Use of Animals for Scientific Purposes (8th Edition, National Health and Medical Research Council, 2013). Male C57BL/6 mice were purchased from Ozgene Animal Resource Centre (Western Australia).

Eight-week-old male C57BL/6J mice were fed a high fat diet (HFD; 60% kcal from fat; SF13-092, Specialty Feeds Pty Ltd, Australia) for 21–24 weeks, as indicated. In the disease-progression model, mice received weekly subcutaneous injections of either control antisense oligonucleotides (control ASO: CCTTCCCTGAAGGTTCCTCC) or a MOE gapmer ASO targeting MLKL (MLKL ASO#2: CGCTAATTTGCAACTGCATC).

In the intervention model, mice were maintained on HFD for 12 weeks to induce obesity, followed by weekly subcutaneous injections of two independent MOE gapmer ASOs targeting MLKL (MLKL ASO#1: ACCCAATTCTTTGTAAGCAA or MLKL ASO#2: CGCTAATTTGCAACTGCATC) or RIPK1 (RIPK1 ASO#1: TCAGCCACTTCTGAAGCATT or RIPK1 ASO#2: CTCCATGTACTCCATCACCA). Antisense oligonucleotides were generously provided by Ionis Pharmaceuticals (USA).

At study termination, mice were fasted for 6 h and anaesthetised with isoflurane prior to cardiac puncture, followed by exsanguination and PBS or saline perfusion, as previously described [15, 32].Tissue samples were either snap-frozen in liquid nitrogen and stored at −80°C or fixed in 10% neutral-buffered formalin (YAMS-NBF; Trajan Scientific and Medical, Australia) and stored at 4°C for subsequent analysis. Frozen adipose tissue was embedded into Optimal Cutting Temperature (OCT) compound and sectioned at 6 μm thickness for histological staining.

### Glucose and insulin tolerance tests

Following a 6 h fast, mice received intraperitoneal injections of either D-glucose (1 g/kg body weight; #1083371000, Merck Australia) for glucose tolerance tests or insulin (0.75 U/kg body weight; Humalog, ARTG 53488, Eli Lilly Australia) for insulin tolerance tests. Tail-vein blood glucose was measured at designated time points using an Accu-Chek Aviva Nano glucometer (Roche). Glucose values were plotted over time, and the area under the curve (AUC) was calculated for each animal. Data are presented as mean ± SEM.

### Energy Expenditure and Indirect Calorimetry

Body composition (fat mass, lean mass, free water, and total body water) was measured using an EchoMRI™ quantitative nuclear magnetic resonance (NMR) body composition analyser, as previously described [15]. Mice were then individually housed in the TSE PhenoMaster system (TSE Systems, Germany) to assess energy expenditure, locomotor activity, and food and water intake. Each plexiglass metabolic chamber was supplied with continuous airflow (0.5 L/min) and maintained at 25°C under a standard 12-h light/dark cycle. Following a 48–60 h acclimation period, metabolic parameters were recorded over an additional 48 h, with data collected hourly from each chamber. Data were analysed using CalR (beta), a web-based analysis tool for indirect calorimetry experiments [33].

### Histopathological Assessment of MASLD

Approximately 20 mg of liver tissue from the left lobe was fixed in 10% neutral-buffered formalin at 4°C, embedded in optimal cutting temperature (OCT) compound, and cryosectioned at 6 μm using a Leica CM1950 cryostat. Sections were stained with haematoxylin and eosin (H&E) and Masson’s trichrome. Blinded histological evaluation was performed according to the Kleiner classification for fibrosis [34] and the NAFLD Activity Score (NAS) [35]. Steatosis was graded as 5–33% (grade 1), >33–66% (grade 2), or >66% (grade 3) of parenchyma involved. Lobular inflammation and hepatocyte ballooning were graded as described previously. Fibrosis staging followed Kleiner criteria. Non-alcoholic steatohepatitis (NASH) was defined by the combined presence of steatosis, lobular inflammation, and hepatocyte ballooning (NAS ≥3), in accordance with European Association for the Study of the Liver (EASL) clinical practice guidelines.

### Serum markers measurement

Serum cytokines were measured using Cytometric Bead Array mouse inflammation kit (#552364, BD Biosciences, USA). In brief, serum samples and standards were prepared according to the manufacturer’s recommendation. Samples and standards were then run through Attune CytPix flow cytometer (Thermo Fisher Scientific, USA). Gating was followed by the instruction manual provided by the company. Other serum markers, including ALT, AST, cholesterol and triglycerides, were analyzed by Monash Pathology using automated AU/DxC Beckman Coulter analyzers.

### Cell Culture and Adipocyte Differentiation

3T3-L1 preadipocytes (#CL-173, ATCC, USA) were cultured in DMEM containing 4.5 g/L glucose, sodium pyruvate, and glutamine, supplemented with 10% newborn calf serum (#16010159, Thermo Fisher Scientific, Australia) on wells coated with 0.1% gelatin. Cells were maintained at 37°C in 5% CO₂ and passaged before reaching 70% confluence. Adipocyte differentiation was induced at confluence as previously described [36]. Briefly, cells were maintained at 100% confluence for 48 h before the addition of differentiation medium containing 0.5 mM isobutyl methylxanthine (IBMX; #I7018, Merck), 0.25 μM dexamethasone (#D1756, Merck), 200nM insulin (#I9278, Merck) and 2 μM Rosiglitazone (#R2408, Merck) for 3 days. Cells were then maintained in culture medium containing insulin alone (200 nM) until days 8–10 post-differentiation, when experiments were performed.

### Generating CRISPR/Cas9 MLKL knockout cells

CRISPR/Cas9 constructs targeting murine *Mlkl* were generated by cloning sgRNAs (antisense: GCACACGGTTTCCTAGACGC or GGCTGCGCACACTCATTGTG; sense: GATGCAGTTGCAAATTAGCG or AGGAACATCTTGGACCTCCG) into the mMLKL-LentiCRISPR v2-mCherry vector, which co-expresses Cas9 and mCherry for fluorescent enrichment as described [37]. Lentiviral particles were produced in HEK293T cells using pCMV-ΔR8.2 (gal/pol) and pMD2.G (VSV-G) helper plasmids. Viral supernatants were collected, filtered (0.45μM), aliquoted and stored at -80°C. For transduction, thawed virus was supplemented with polybrene (5μg/mL) and added to 3T3-L1 adipocytes. Doxycycline (1μg/mL) was included where required for construct induction. For clonal isolation, cells were trypsinised and sorted on a BD Aria^TM^ Fusion Flow Cytometer (sterile mode; 100μM nozzle; 561nM excitation) to isolate single mCherry-positive and fluorescence-negative cells into 96-well plates for expansion.

Genomic DNA from clonal lines was used for amplicon sequencing to verify on-target editing. The *Mlkl* locus was amplified using the NGS forward primer sets listed below (Table 1) and clones containing biallelic frameshift mutations were advanced. Loss of MLKL protein expression was confirmed by western blotting (MLKL normalized to vinculin or total protein).

**Table 1:**
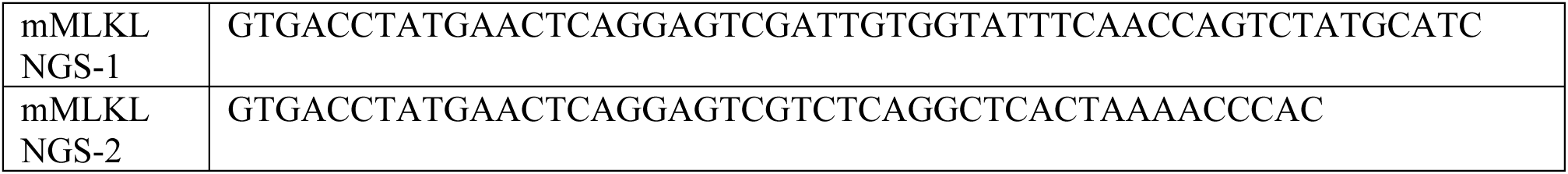
NGS Forward Primer Sets.

### siRNA Transfection

On day 5 of differentiation, 3T3-L1 adipocytes were transfected with ON-TARGETplus mouse MLKL siRNA (L-061420-00-0010, Millennium Science, Australia) or scrambled control siRNA (DHA-D-001810-10-20, Millennium Science, Australia) using Lipofectamine 3000. Cells were exposed to 250 nM siRNA for 4 h, after which the medium was replaced to achieve a final siRNA concentration of 50 nM. Experiments were performed 72 h post-transfection. Knockdown efficiency was confirmed by western blotting as previously described [38].

### Glucose Uptake Assay

Glucose uptake was assessed using the fluorescent glucose analogue 2-NBDG [2-[N-(7-nitrobenz-2-oxa-1,3-diazol-4-yl)amino]-2-deoxyglucose; # N13195, Thermo Fisher Scientific). 24 h-prior to the assay, cells were incubated in DMEM containing 0.5% Fetal Bovine Serum (#FBSFR-S00JF, Scientifix, Australia) and treated with recombinant TNF (50 ng/mL), palmitic acid (500 μM in 1% fatty acid-free BSA), or vehicle control (fatty acid-free BSA). On the day of the experiment, cells were serum- and glucose-starved for 1 h, followed by stimulation with or without insulin (100 nM) for 2 h. 2-NBDG (100 μM) was added for 30 min, after which cells were washed with ice-cold PBS and lysed in 0.1% Triton X-100. Fluorescence was measured (Ex 465 nm/Em 540 nm) using a VantaStar microplate reader (BMG Labtech, Germany) and normalised to control conditions. Glucose uptake was expressed as relative fluorescence units (RFU) normalized to control conditions.

### Quantitation of Lipid Droplets or total lipids

Differentiated adipocytes were treated with vehicle BSA control or palmitic acid for 72 h and stained with 2 μM BODIPY 493/503 9(#D3821, Thermofisher) and 2 μg/mL Hoechst 33342 (#HB0787, Genesearch) for 30 min at 37°C. Cells were washed and either imaged live or fixed in 4% formaldehyde. Images were acquired using a Nikon ECLIPSE Ts2-FL microscope (10× magnification) and analysed using ImageJ. Lipid droplet size and number were quantified using automated particle analysis with predefined size (0–∞) and circularity (0.5–1.0) thresholds as previously indicated [39]. Prior to quantification, images were processed using standard filtering and watershed algorithms to separate overlapping lipid droplets. For total lipid content analysis, cells were lysed in 0.1% Triton X-100, and fluorescence was measured using a microplate reader (VantaStar, BMG Labtech, Germany) at an excitation wavelength of 465 nm and emission wavelength of 540 nm. Lipid content was expressed as relative fluorescence units (RFU) normalised to control conditions.

### Glycerol Release Assay

Lipolysis was assessed by measuring glycerol release from differentiated adipocytes. Cells were serum-starved in culture medium containing 0.5% fetal bovine serum (FBS) for 24 h and then treated with the β-adrenergic receptor agonist isoprenaline (1 μM; #I2760, Merck) for 2 h. Culture medium was collected, and glycerol concentration was measured using a colorimetric assay (#MAK117, Sigma-Aldrich) according to the manufacturer’s instructions. Lipolytic activity was expressed as glycerol release per mg protein.

### Western Blotting

Cells were treated overnight with TNF, palmitic acid, or vehicle control, followed by stimulation with insulin (100 nM) for 1 h. Cells were lysed in NuPAGE LDS sample buffer (#NP0007, Thermo Fisher Scientific) supplemented with 10% β-mercaptoethanol (#M6250, Sigma-Aldrich). Proteins were separated by SDS-PAGE and transferred to PVDF membranes.

Membranes were blocked and incubated overnight at 4°C with primary antibodies diluted 1:2000 in PBS against phospho-AKT (Ser473; #9271, Cell Signaling Technology), total AKT (#9272, CST), MLKL (clone 3H1; MABC604, Merck), and vinculin (#13901, CST). Membranes were then incubated with HRP-conjugated secondary antibodies (anti-rabbit #7074 or anti-rat #7077; CST). Proteins were visualised using enhanced chemiluminescence and imaged on a ChemiDoc XRS+ system (Bio-Rad, USA).

### RT-qPCR

Tissues and treated cells were lysed using the lysis buffer supplied with the ISOLATE II RNA Mini Kit (52073, Meridian Bioscience). Liver and adipose tissues were homogenised in lysis buffer using a bullet blender. Total RNA was extracted according to the manufacturer’s instructions. For cDNA synthesis, 500 ng of total RNA was reverse-transcribed using the iScript™ cDNA Synthesis Kit (1708891, Bio-Rad). For SYBR Green–based qPCR, reactions were performed using PowerTrack™ SYBR Green Master Mix with 5 ng/μL cDNA input and manufacturer-recommended cycling conditions. Threshold cycle (Ct) values were obtained, and mRNA expression levels were calculated by gene copy number and normalised to the housekeeping gene as indicated (e.g. *Ppia*). For TaqMan-based qPCR, reactions were performed using TaqMan™ Fast Universal PCR Master Mix (#4352042, Thermo Fisher Scientific) with *Ripk1* (Mm00436354_m1)*, Mlkl* (Mm01244222_m1) or *Hprt* (Mm03024075_m1) probes and 5 ng/μL cDNA input. Amplification was carried out using standard TaqMan cycling conditions on a real-time PCR system. Relative gene expression was calculated using the ΔΔCt method and normalised to *Hprt*. Primer and probe details are provided below.

**Table 2:**
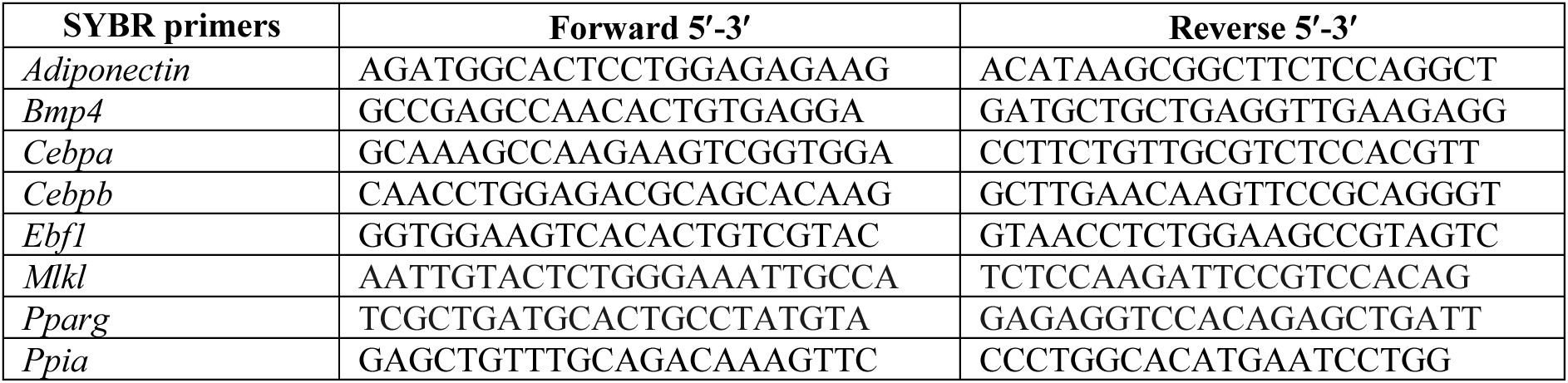
SYBR Primers.

### Statistical Analysis

Data are presented as mean ± standard error of the mean (SEM) from at least three independent experiments performed in triplicate, or from n=8-12 mice per group, as indicated. Statistical analyses were performed using GraphPad Prism (version 10, GraphPad Software, San Diego, CA, USA). Comparisons between two groups were performed using unpaired two-tailed Student’s t-test. For comparisons involving more than two groups, one- or two-way analysis of variance (ANOVA) was used, followed by appropriate post-hoc multiple-comparison tests, as specified. Statistical significance was defined as p < 0.05.

## RESULTS

### MLKL suppression reduces adiposity in obese mice and improves insulin sensitivity

We investigated the hypothesis that knockdown of mixed lineage kinase domain-like protein (MLKL), previously shown to be protective in other cardiometabolic conditions [20, 21, 40, 41], may also attenuate diet-induced obesity and insulin resistance. To assess this, we examined whether long-term MLKL silencing using antisense oligonucleotides (ASOs) could ameliorate metabolic dysfunction in a mouse model of high fat diet (HFD)-induced obesity. Male C57BL/6J mice were fed a HFD (60% kcal from fat) for 24 weeks while simultaneously receiving weekly injections of either a control ASO or one of two MLKL-targeting ASOs (MLKL ASO#2, 50mg/kg). MLKL ASO treatment significantly reduced whole-body weight during HFD feeding (**Figure 1A**). This reduction in body weight was primarily attributable to a marked decrease in total fat mass, with no detectable changes in lean mass, as determined by EchoMRI after 16 weeks of HFD (15.88 ± 4.86 in control vs 6.96 ± 4.11 grams in MLKL ASO#2; p=0.000075; **Figure 1B**). Consistent with these findings, epididymal and subcutaneous fat pad weights, as well as liver weight, were significantly reduced in MLKL-ASO#2-treated mice at the study endpoint, with spleen weight unaffected (**Figure 1C**; p<0.05). We have previously demonstrated effective delivery of 2**′**F-MOE ASOs to liver and adipose tissue [15], and accordingly, *Mlkl* mRNA expression was markedly reduced in these tissues (**Figure 1D**). In agreement with previously published work [40, 42, 43], MLKL inhibition reduced hepatic steatosis but did not significantly alter hepatic inflammation, ballooning or fibrosis (**Figures 1E-F**; **Supplemental Figure 1A**). Furthermore, similar to observations with RIPK1 ASO therapy [15], adipocyte size in both subcutaneous and epididymal depots was markedly reduced in MLKL ASO#2–treated mice (**Figures 1G–H; Supplemental Figures 1B–C**). This reduction in adipocyte size is indicative of potential decreased triglyceride storage, consistent with reduced circulating triglyceride levels (**Figure 1I**), while total circulating cholesterol levels remained unchanged (**Supplemental Figure 1D**). Importantly, MLKL ASO therapy did not alter serum liver enzyme levels (AST or ALT; **Supplemental Figure 1E**), nor did it induce expression of interferon-stimulated genes (*Ifit1, Ifit3, Oasl, Mx1*; **Supplemental Figure 1F**), a concern raised in prior ASO-based therapies [44], further supporting the overall safety and tolerability of this ASO-knockdown approach. Moreover, serum levels of key cytokines associated with pro- or anti-inflammatory responses did not differ between groups, suggesting that MLKL inhibition primarily modulates the local adipose tissue microenvironment rather than systemic inflammation (**Supplemental Figure 1G**).

**Figure 1:**
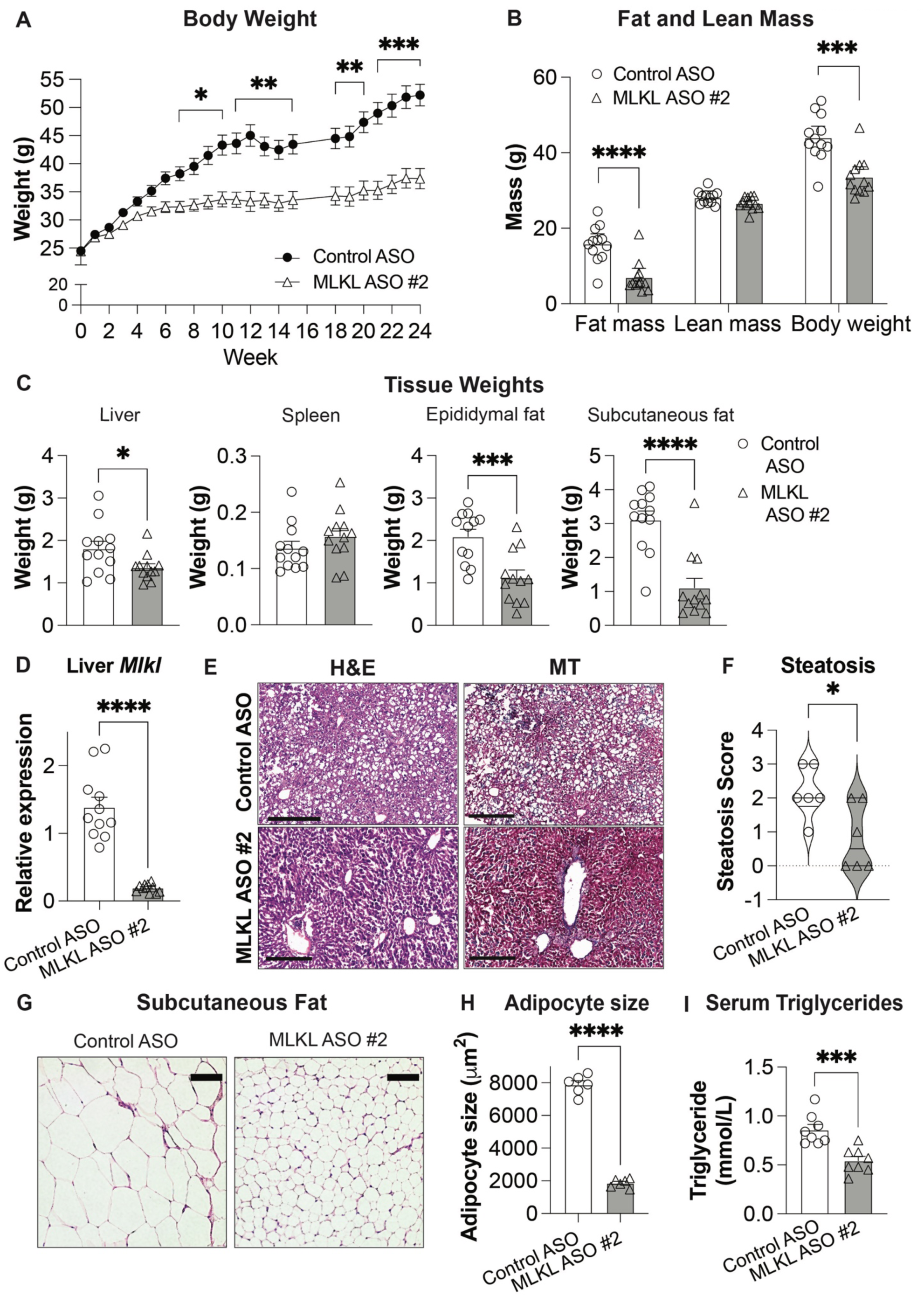
Therapeutic knockdown of MLKL in a diet-induced obesity model protects against obesity. Male C57BL/6J mice (n=12/group) were fed a high-fat diet (60% kCal) for 24 weeks with weekly injections of 50mg/kg control ASO or MLKL ASO#2. (**A**) Body weight. Data are mean ± SEM and were analysed by one-way ANOVA with Dunnett’s multiple-comparison test; *p < 0.05, **p < 0.01 and ***p<0.001. (**B**) Fat, lean and total body mass measured by EchoMRI (NMR). (**C**) Liver, Spleen, epididymal and subcutaneous adipose tissue weight at termination of the study. (**D**) Hepatic *Mlkl* mRNA expression after 24 weeks. (**E**) Representative liver haematoxylin and eosin (H&E) stain and Masson’s trichrome (MT) stain. (**F**) Liver steatosis scoring. (**G**) Representative subcutaneous adipose tissue H&E images. (**H**) Adipocyte size quantification. Representative epididymal fat and respective adipocyte size is shown in Supplemental Figure 1 (E& F). (**I**) Fasting serum triglycerides. (**F, H**) Violin plot and bar graph depict the mean ± SEM of an average of five sections per mouse (n=8 mice/group). (**B-I**) Significance assessed using a two-tailed Student’s *t*-test; *p < 0.05, ***p<0.001, and ****p<0.0001.

Given the substantial reductions in total fat mass and white adipose tissue depots (epididymal and subcutaneous), we next assessed whole-body metabolic function using an open-circuit indirect gas calorimetry system to quantify oxygen consumption and carbon dioxide production. MLKL ASO#2–treated mice exhibited a significantly lower respiratory exchange ratio (RER) over the 48-hour monitoring period compared with control mice (**Figure 2A–B**), despite equivalent food intake between groups (**Supplemental Figure 2A**). This reduction in RER suggests enhanced metabolic flexibility, characterized by a greater reliance on fatty acid oxidation for energy utilization. Consistent with these metabolic changes, MLKL ASO#2–treated mice displayed increased locomotor activity during the nocturnal (dark) 12-hour phase relative to controls (**Figure 2C–D**), indicating preserved circadian rhythmicity and increased energy expenditure during active phase. Furthermore, consistent with metabolic improvements observed with RIPK1 ASO therapy [15], MLKL ASO#2 treatment also enhanced glucose homeostasis, as evidenced by improved glucose control at 14 weeks and improved insulin sensitivity at 19 weeks of high-fat diet feeding [glucose tolerance test (GTT), **Figure 2E**; insulin tolerance test (ITT), **Figure 2F**] at 14 weeks and 19 weeks HFD-feeding respectively. Together, these findings show that long-term MLKL silencing through ASO therapy confers broad metabolic benefits in diet-induced obese mice, including substantial reductions in adiposity, enhanced metabolic flexibility and improved glucose homeostasis, highlighting MLKL as a promising therapeutic target for mitigating obesity-associated metabolic dysfunction.

**Figure 2:**
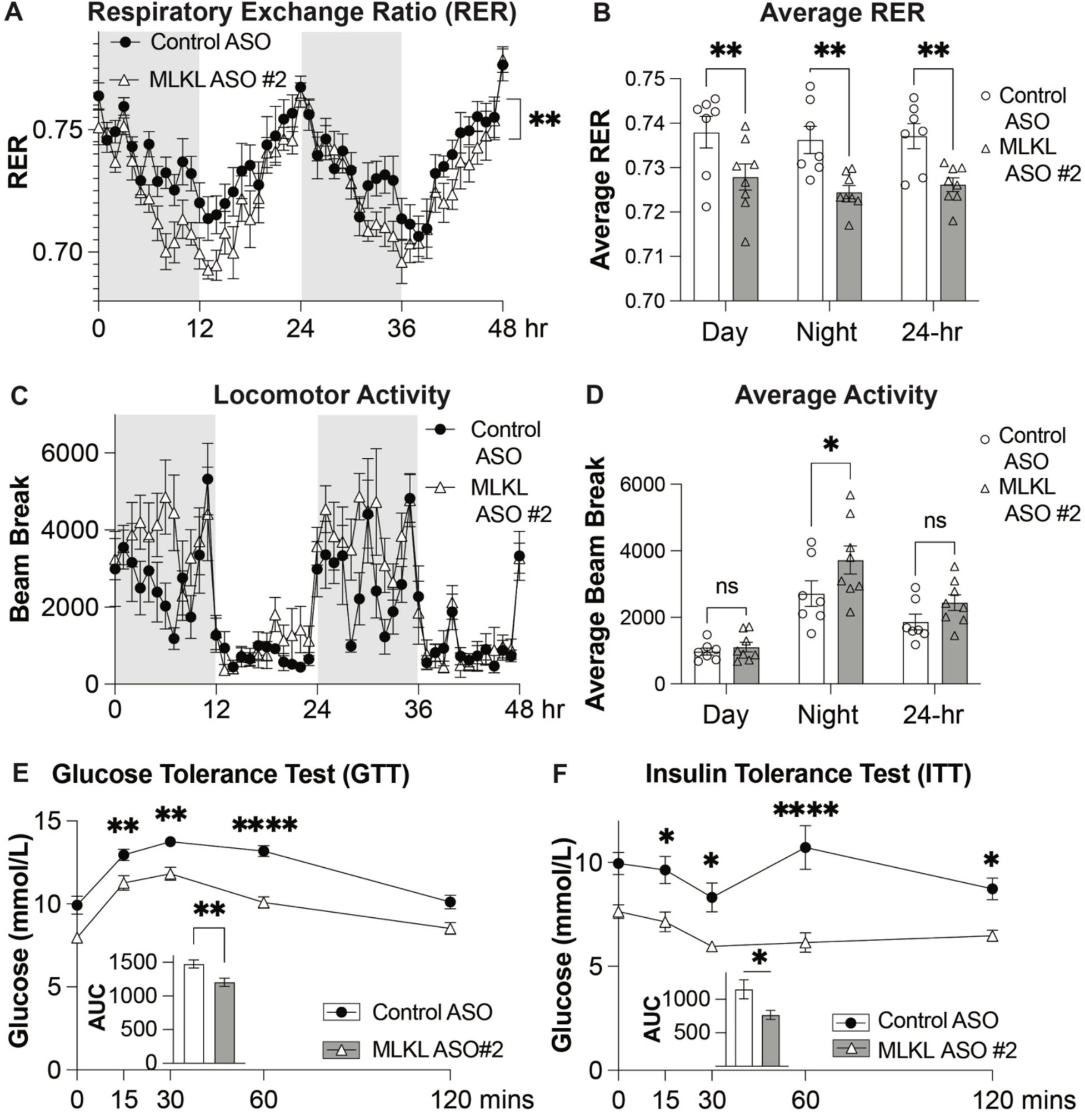
MLKL knockdown promotes fat metabolism, activity and improves glucose homeostasis. Between weeks 16 and 17 of ASO-treatment, mice (n=8/group) were individually housed in the TSE Phenomaster indirect calorimetry system to assess metabolic parameters. (**A**) Respiratory exchange ratio (RER=VCO_2_/O_2_) measured over 48 h. (**B**) Average RER during 12-h light (day), dark (night), and 24-h periods. (**C**) Locomotion activity quantified as beam breaks per hour over 48 h. **(D)** Average locomotor activity during day, night and 24 h periods. Food consumption over 48 h is shown in Supplemental Figure 2. (**A, C**) Grey shading denotes the dark (night) phase and white shading denotes the light phase. Data were analysed using CalR beta software. (**B, D**) Bar graphs show mean ± SEM; significance was assessed using two-way ANOVA with Tukey’s multiple comparison test; *p<0.05, **p<0.01. (**E**) Glucose tolerance test (GTT) and (**F**) Insulin tolerance test (ITT) performed after 12 or 18 weeks of treatment respectively; AUC, area under curve. Data are mean ± SEM (n=12/ group) and were analysed using two-way ANOVA and Tukey’s multiple-comparison test; *p < 0.05, **p < 0.01 and ****p < 0.0001.

### Targeted MLKL knockdown in mature adipocytes reduces lipid accrual and enhances lipolysis

To define the cell-autonomous role of MLKL in adipocytes, we generated CRISPR/Cas9 MLKL-knockout (MLKL-KO) 3T3-L1 cells as described [37] and verified loss of MLKL protein via western blotting (**Figure 3A, C**). Under basal conditions and following palmitate challenge, MLKL-KO adipocytes accumulated less neutral lipid than wild-type (WT) cells, as shown by BODIPY fluorescence imaging (**Figure 3B**) and fluorometric quantification across a palmitate titration (**Figure 3D**), indicating a reduced ability for MLKL-KO cells to accumulate lipid. We therefore assessed genes associated with adipogenic programming. During differentiation (day 0 to 8; D0-D8), WT 3T3-L1 cells exhibited the expected progressive induction of *Adipoq*, *Cebpa* and *Pparg* over the experimental period up to 8 days, with *Cebpb* peaking at Day 2 and then declining thereafter (**Figure 3E**, p<0.0001). By contrast, MLKL KO 3T3-L1 cells show blunted induction of *Adipoq*, *Cebpa* and *Pparψ*, with only a modest reduction in *Cebpb* (**Figure 3E**, p<0.05). Collectively, these data indicate that MLKL-KO cells lose their adipogenic identity, potentially due to impairment of the lipid-filling phase of adipogenesis.

**Figure 3.**
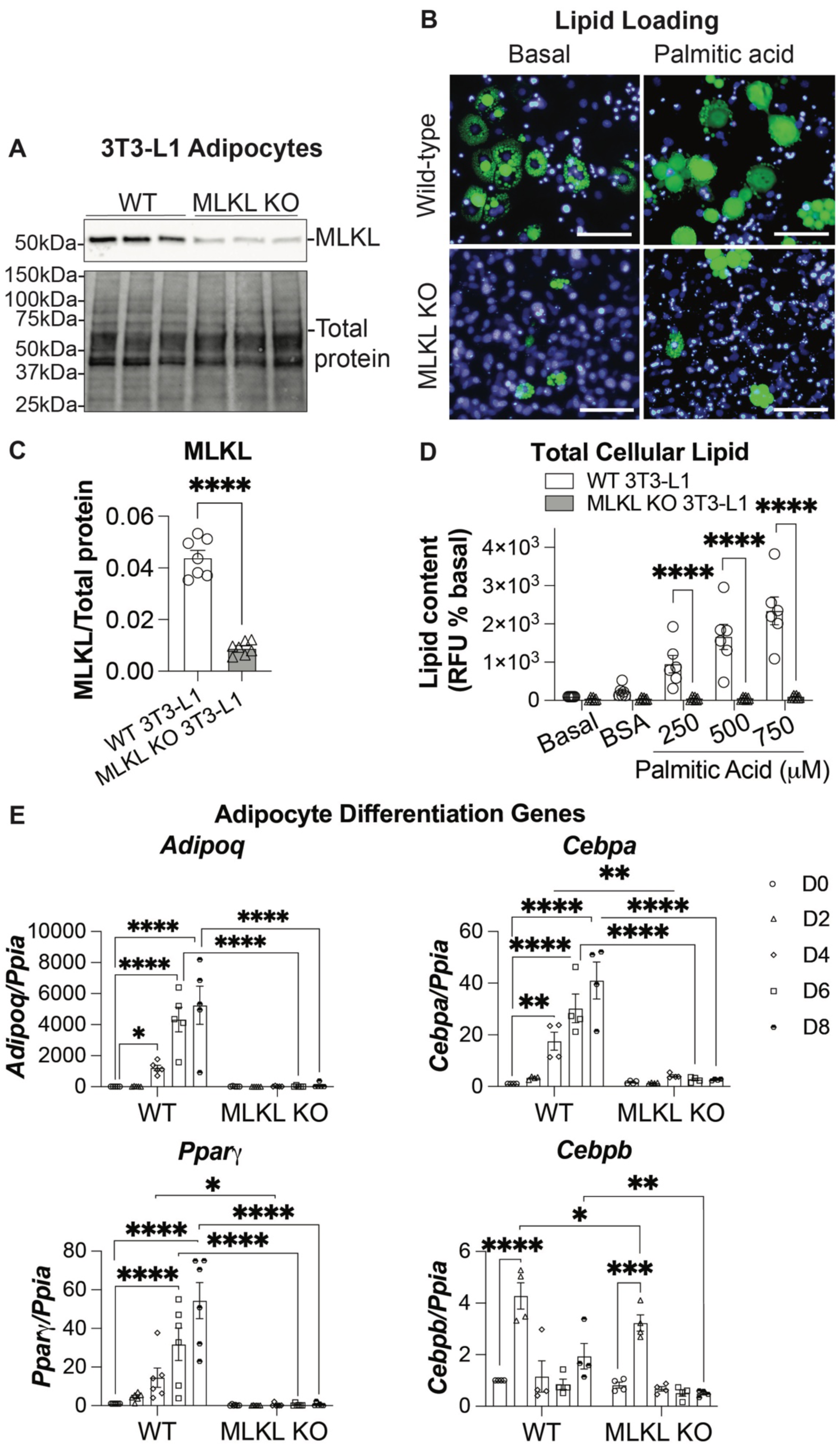
Loss of MLKL impairs lipid accumulation and adipocyte differentiation in 3T3-L1 adipocytes. (**A**) Representative western blot confirming MLKL deletion in CRISPR-generated MLKL knockout (KO) compared to wild-type (WT) 3T3-L1 adipocytes; total protein staining served as a loading control. (**B**) Representative fluorescence microscopy images of BODIPY-labelled lipid droplets (green) and Hoechst-stained nuclei (purple) in WT and MLKL KO adipocytes under basal conditions or treatment with palmitic acid (750 µM). Scale bars=100 µm. **(C)** Quantification of MLKL protein expression normalised to total protein (n=7). **(D)** Total cellular lipid content assessed by BODIPY fluorescence (relative fluorescence units, RFU, % basal) in WT and MLKL KO adipocytes treated with BSA vehicle (basal) or palmitic acid (250, 500, or 750 µM) (n=6). **(E)** Relative mRNA expression of adipogenic markers (*Adipoq*, *Cebpa*, *Pparγ*, and *Cebpb*) normalised to housekeeping gene, *Ppia* across the differentiation time course (day 0 to 8; D0–D8) in WT and MLKL KO cells (n=4-6). Data are mean ± SEM. Statistical significance was determined by two-way ANOVA with Tukey’s multiple comparisons test. *p<0.05, **p<0.01, ***p<0.001, ****p<0.0001.

Since post-development MLKL knockdown in mice with ASOs increased fat oxidation (**Figure 2A-B**), we next asked whether post-differentiation MLKL knockdown in mature 3T3-L1 adipocytes with siRNAs would similarly promote lipid mobilisation. In day-5 differentiated 3T3-L1 adipocytes – corresponding to the active lipid-filling phase of adipogenesis. – siRNA-mediated MLKL knockdown was confirmed by western blotting (**Figure 4A**). Upon palmitate exposure, MLKL-silenced cells displayed smaller lipid droplets and reduced total cellular lipid compared with control siRNA cells, as shown by BODIPY fluorescence imaging (**Figure 4B)** and by quantification of droplet size and bulk lipid content across increasing palmitate concentrations (**Figure 4C–D**). Given the reduction in stored lipid, we assessed lipolysis. In the presence of palmitate, MLKL-siRNA cells released more glycerol than controls (**Figure 4E**). Furthermore, β-adrenergic stimulation with isoprenaline increased glycerol release under both vehicle (BSA) and agonist conditions, with consistently greater responses in MLKL-silenced cells (**Figure 4F**). These findings suggest that, rather than complete MLKL deletion in preadipocytes, targeted MLKL knockdown in mature adipocytes more selectively limits lipid deposition and enhances lipid mobilization, supporting a potentially more tractable therapeutic strategy.

**Figure 4.**
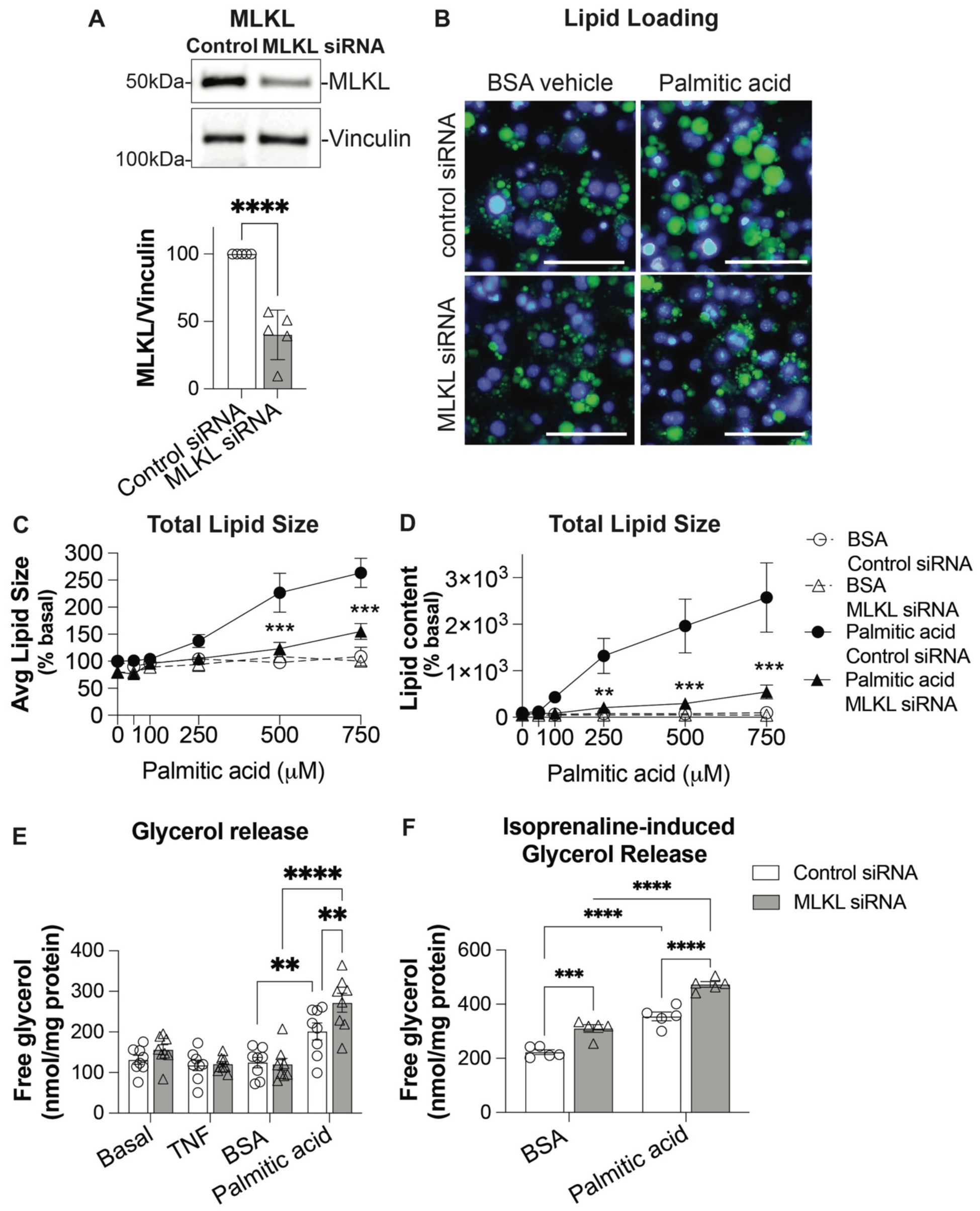
MLKL knockdown reduces lipid loading and enhances lipolysis in 3T3-L1 adipocytes. 3T3-L1 adipocytes were transfected with control or MLKL siRNA (50nM) at day 5 post-adipogenesis, and cells were analysed on day 8. **(A)** Representative western blot of MLKL protein expression, vinculin as a loading control and quantification of MLKL expression normalised to vinculin (n=5). **(B)** Representative fluorescence microscopy images of BODIPY-stained lipid droplets (green) and Hoechst-stained nuclei (purple) under basal (BSA control) conditions or palmitic acid treatment (750 µM). Scale bars = 100 µm. **(C)** Average lipid droplet size (% basal) in response to increasing concentrations of palmitic acid (0–750 µM) or fatty acid-free BSA vehicle (n = 5). **(D)** Total cellular lipid content (% basal) following palmitic acid or BSA treatment (n = 4). **(E)** Basal lipolysis assessed by free glycerol release (nmol/mg protein) under basal conditions or following treatment with TNF (50 ng/ml), BSA vehicle, or palmitic acid (750 µM) (n = 8). **(F)** Isoprenaline (1 µM)-stimulated glycerol release in cells treated with BSA vehicle or palmitic acid (750 µM) (n = 5). Data are mean ± SEM. Statistical significance was determined using two-way ANOVA with Tukey’s multiple-comparisons test; *p < 0.01, **p < 0.001, ***p < 0.0001.

### MLKL knockdown preserves glucose homeostasis during inflammatory or metabolic stress

Insulin activates its receptor to trigger a signalling cascade culminating in AKT phosphorylation, a key step required for glucose uptake, lipid metabolism, and overall metabolic homeostasis. Building on the *in vivo* improvement in whole-body glucose control, we next tested whether post-differentiation MLKL silencing enhances insulin-AKT signalling glucose uptake in mature adipocytes exposed to inflammatory or metabolic stress. In mature 3T3-L1 adipocytes, MLKL siRNA maintained robust insulin-stimulated AKT phosphorylation (p-AKT/AKT) relative to control siRNA (**Figure 5A**). Under non-stressed conditions, insulin increased AKT phosphorylation to a similar extent in both groups. However, TNF markedly reduced insulin-induced phosphorylation of AKT in control adipocytes, whereas MLKL silencing preserved the insulin-AKT response (**Figure 5A, left**). Similarly, palmitic acid blunted insulin-AKT signalling in control cells but MLKL knockdown restored AKT phosphorylation to vehicle-control levels (**Figure 5A, right**). Consistent with improved proximal signalling, MLKL-siRNA treated mature adipocytes displayed higher insulin-stimulated glucose uptake than control cells (**Figure 5B**). Notably, MLKL knockdown mitigated the inhibitory effects of TNF and palmitate on glucose uptake, restoring responses toward unstressed levels (**Figure 5B**). Taken together, with our *in vivo* data, these cell-autonomous findings support a model in which MLKL suppression shifts mature adipocytes toward increased lipid mobilisation and fat oxidation, while preserving insulin–AKT signalling and glucose uptake, thereby contributing to improved metabolic health and reduced obesity (**Figure 5C**).

**Figure 5.**
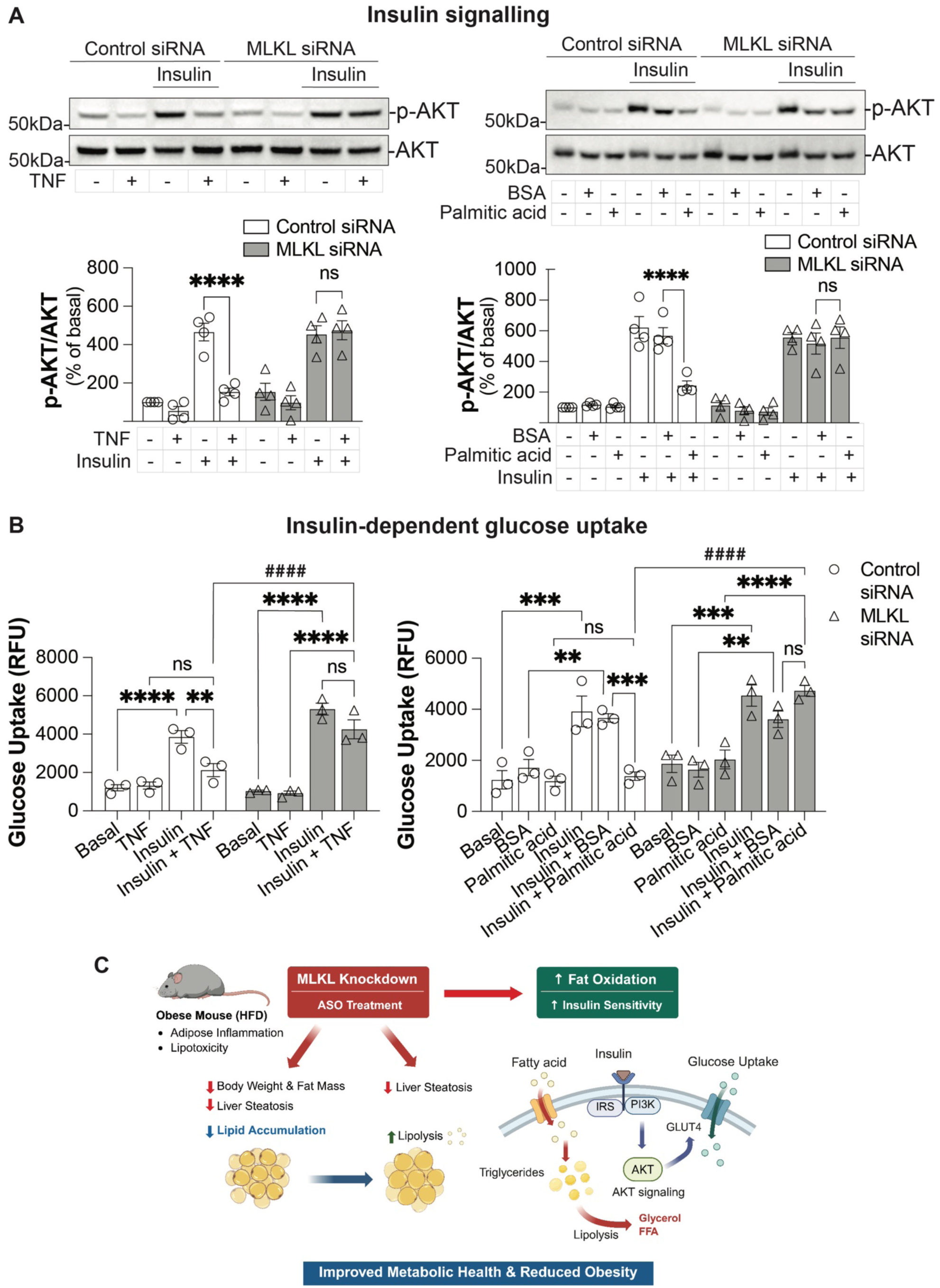
MLKL knockdown rescues lipotoxic- and inflammatory stress–induced impairment of insulin signalling and glucose uptake. **(A)** Representative western blots and quantification of insulin-stimulated AKT phosphorylation (Ser473; p-AKT), expressed as p-AKT/total AKT (% basal), in control or MLKL siRNA treated 3T3L-1 adipocytes stimulated with insulin (100 nM) in the presence or absence of TNF (50 ng/ml) (left panels; n=4), or fatty acid-free BSA vehicle and palmitic acid (500 µM) (right panels; n=4). **(B)** Insulin-dependent glucose uptake (Relative Fluorescence Units, RFU) in control and MLKL siRNA-transfected 3T3-L1 adipocytes under basal conditions or following treatment with TNF (50 ng/ml) and/or insulin (100 nM) (left panel; n=3), or BSA vehicle or palmitic acid (750 µM) with or without insulin (100 nM) (right panel; n=3). Data are mean ± SEM. Statistical significance was determined by two-way ANOVA with Tukey’s multiple comparisons test. **p<0.01, ***p<0.001, ****p<0.0001; ^####^p<0.0001 indicates a significant difference in insulin-stimulated glucose uptake between control and MLKL siRNA groups in the presence of TNF or palmitic acid. **(C)** Schematic summary of metabolic effects of MLKL knockdown in adipocytes. Proposed model illustrating that MLKL knockdown reduces lipid accumulation and enhances lipolysis and fatty acid oxidation, thereby preserving insulin signalling and glucose uptake under lipotoxic and inflammatory conditions. These effects contribute to improved metabolic health and reduced obesity-associated insulin resistance. Created with Biorender.com.

### Therapeutic intervention targeting RIPK1 or MLKL reverses metabolic complications

Leveraging the adipocyte-intrinsic effects of MLKL knockdown, we investigated whether therapeutic intervention with RIPK1 or MLKL ASOs in established obesity attenuates adiposity and restores insulin sensitivity. Male C57BL/6J mice were fed a high-fat diet for 12 weeks to induce obesity, then treated weekly with two independently targeting 2′MOE gapmer ASOs against RIPK1 (ASO#1 or ASO#2; 50 mg/kg) or MLKL (ASO#1 or ASO#2; 50 mg/kg) until study termination at weeks 21–24. As in the progression study [15], no differences were observed between no-treatment (NT) and control ASO groups (**Supplemental Figure 3A**), indicating that control ASO did not measurably affect metabolic phenotypes assessed and serves as an appropriate control.

RIPK1 or MLKL ASO therapy efficiently suppressed target transcripts in liver and adipose tissue – *Ripk1* in RIPK1 ASOs (**Figure 6A; Supplemental Figure 3B**) and *Mlkl* with MLKL ASOs (**Figure 6D; Supplemental Figure 3C**), consistent with effective delivery. In obese mice receiving RIPK1 ASO#1 or #2, the progressive weight gain observed with control ASO was halted from ∼week 14 onward and body weight thereafter remained lower (**Figure 6B**). EchoMRI showed that fat-mass reduction accounted for the majority of weight change, with lean mass preserved (**Figure 6C**, p<0.05). Similarly, in mice treated with MLKL ASO#1 or #2, body weight declined by week 18, again driven predominantly by loss of fat mass (**Figure 6E**; p<0.0001), with lean mass unchanged (**Figure 6F**). Metabolic testing demonstrated improved glucose tolerance in both intervention arms versus control ASO (**Figure 6G-H**). Together with our *in vitro* findings, these data show that post-obesity intervention targeting RIPK1 or MLKL confers coordinated benefits across liver and adipose tissue, reducing adiposity and improving systemic glucose homeostasis, even when treatment is initiated after obesity is established.

**Figure 6:**
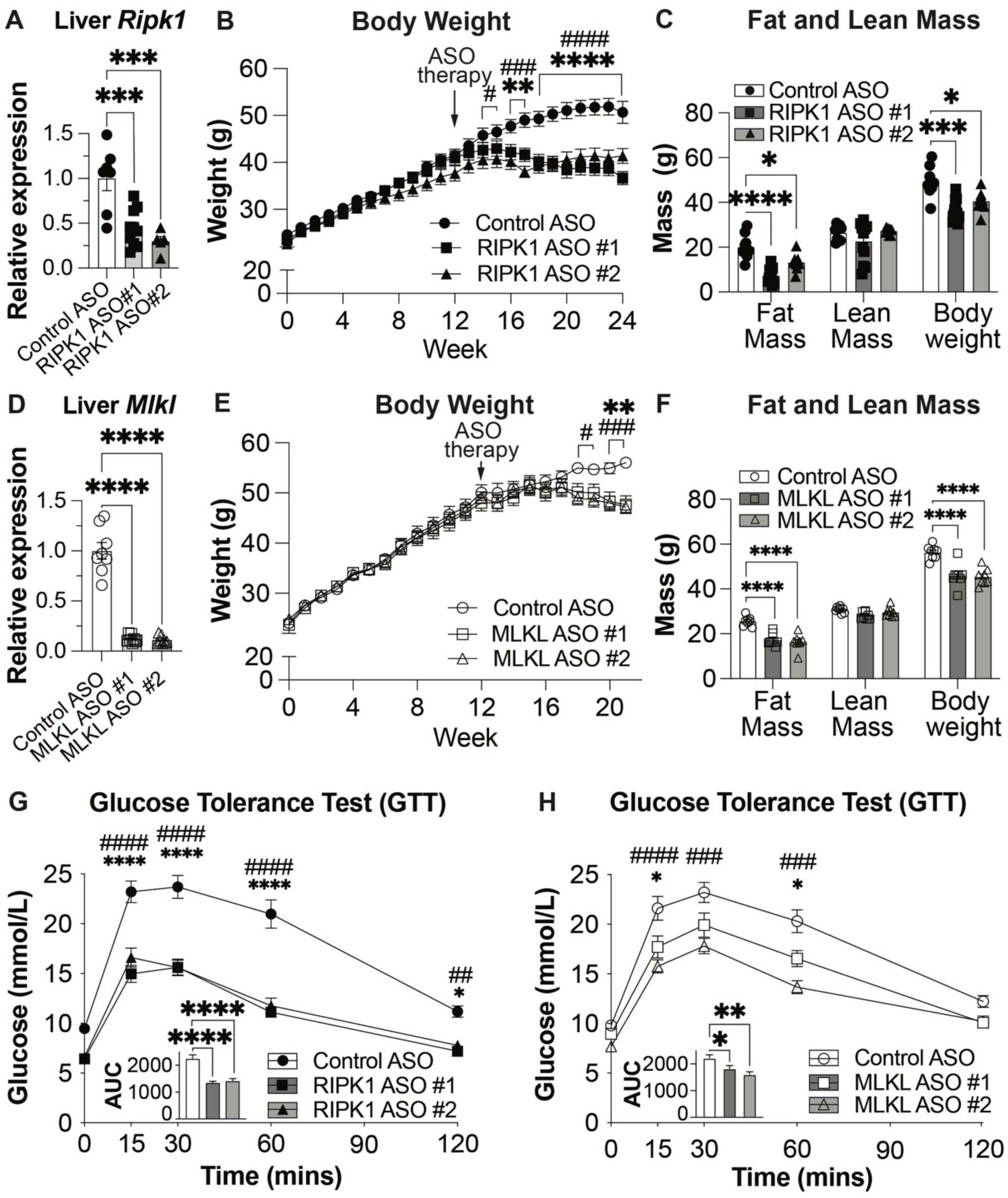
Therapeutic knockdown of RIPK1 or MLKL reverses obesity and insulin resistance in established diet-induced obesity. Male C57BL/6J mice (n=8/group) were fed a high-fat diet for 12 weeks to establish obesity, followed by weekly administration of control ASO, RIPK1 ASOs (#1 or #2) or MLKL ASO (#1 or #2) as indicated. **(A)** Hepatic *Ripk1* mRNA expression following control or RIPK1 ASOs treatment. **(B)** Body weight trajectory during RIPK1 ASO therapy. **(C)** Fat mass, lean mass and body weight following RIPK1 ASOs treatment. **(D)** Hepatic *Mlkl* mRNA expression following MLKL ASOs treatment; adipose *Ripk1* or *Mlkl* expression from both models are shown in Supplemental Figure 3. **(E)** Body weight trajectory during MLKL ASO therapy; comparison between PBS no treatment (NT) or control ASO groups is presented in Supplementary Figure 3A. **(F)** Fat mass, lean mass and body weight following MLKL ASO treatment. **(G, H)** Glucose Tolerance Test (GTT) following RIPK1 ASOs **(G)** or MLKL ASOs **(H)** treatment, with corresponding area under the curve (AUC) shown within the graph. Data are mean ± SEM (n=8 mice/group). For panels **A**, **C**, **D**, **F**, and **G–H** (AUC), statistical significance was assessed using one-way ANOVA with Dunnett’s multiple-comparison test; *p < 0.05, **p < 0.01, ***p < 0.001, ****p < 0.0001. For panels **B**, **E**, and **G–H** (time course), significance was determined using two-way ANOVA with Tukey’s multiple-comparison test. *p < 0.01 and ***p < 0.0001 denote comparisons between control ASO and RIPK1 ASO#1 (**B**) or MLKL ASO#1 (**E**), while ^#^p < 0.05, ^###^p < 0.001, and ^####^p < 0.0001 denote comparisons between control ASO and RIPK1 ASO#2 (**B**) or MLKL ASO#2 (**E**).

## DISCUSSION

In this study, we show that therapeutic MLKL knockdown using two-specific antisense oligonucleotides (ASOs) confers broad metabolic benefits in diet-induced obese mice, reducing total and depot-specific fat mass without loss of lean mass, smaller adipocytes, lower circulating triglycerides, improved glucose tolerance. These effects were accompanied by a lower respiratory exchange ratio (RER) consistent with increased fatty acid oxidation, all occurring without changes in food intake but with increased nocturnal activity. Our findings are consistent with an earlier report in which adipocyte-specific *Mlkl* deletion protected against HFD-induced obesity, enhanced insulin sensitivity, and increased energy expenditure independent of food intake or locomotion, with transcriptional enrichment of oxidative phosphorylation and mitochondrial pathways in white adipose tissue [23]. This suggests adipocyte-intrinsic MLKL promotes triglyceride storage and constrains oxidative metabolism, while MLKL suppression shifts adipocytes toward a more oxidative, lipid-mobilising state that supports whole-body glucose control. Importantly, the metabolic benefits observed with MLKL ASO treatment were phenocopied by ASO-mediated RIPK1 suppression, providing additional evidence that this axis functions through canonical necroptosis signalling rather than through proposed necroptosis-independent roles of MLKL in other physiological processes. Collectively, these findings are broadly concordant with studies reporting that MLKL contributes to diet-induced obesity or metabolic liver disease [17-19, 23]; however, other reports have found minimal or no differences in body weight or insulin resistance in MLKL-deficient mice under similar dietary conditions [22], highlighting context-dependent metabolic roles.

The necroptosis machinery – RIPK1, RIPK3, and MLKL – exerts functions beyond lytic cell death, including key roles in metabolic regulation [17-19, 45, 46]. Our prior work identified RIPK1 as an inflammatory–metabolic node in obesity, with human cis-eQTLs linking higher adipose RIPK1 expression to increased obesity risk. Mechanistically, RIPK1 drives NF-κB–mediated adipose inflammation and insulin resistance in diet-induced obese mice [15]. RIPK1 silencing reduces fat mass and improves insulin sensitivity with favourable adipose immune remodelling, while pharmacological RIPK1 inhibition attenuates steatosis through MLKL-dependent enhancement of hepatic mitochondrial respiration [15, 41, 47]. Consistent with this, RIPK1 targeting ASOs in our intervention study reversed diet-induced weight gain and restored glucose tolerance. In contrast, RIPK3 deletion increases caspase 8–dependent adipocyte apoptosis, amplifies white adipose inflammation, impairs adipose insulin signalling, and causes glucose intolerance [48]. RIPK3 is overexpressed in human visceral fat and correlates with BMI and metabolic markers [48]. Comparative studies therefore distinguish RIPK3–caspase 8 signalling as a driver of macrophage-mediated adipose and hepatic inflammation, whereas MLKL influences obesity-related traits largely independently of inflammation, underscoring non overlapping functions within this axis [18]. Importantly, tissue context and dosage appear critical. Although MLKL mediates necroptosis-associated injury in diabetic heart and kidney [20, 21, 49, 50], multiple studies, including the present work, demonstrates metabolic benefits of MLKL suppression in the absence of overt cell death. Complete MLKL loss can exacerbate hepatic inflammation in some settings, whereas partial reduction, as achieved with ASOs here, improves adiposity and systemic metabolic control [19], favouring a model in which MLKL regulates lipid handling and insulin signalling independently of its canonical membrane-disruptive function. MLKL is also upregulated in human atherosclerosis associated with type 2 diabetes and cardiovascular risk [51], and its loss reduces features of plaque vulnerability despite complex effects on macrophage lipid accumulation [41]. Direct evidence linking MLKL to human adiposity remains limited, but MLKL activation during *in vitro* human stem cell–derived adipogenesis suggests a potential metabolic role [24]. Further, elevated testosterone has been observed in MLKL-deficient male mice [46], an effect hypothesised to reflect altered lipid metabolism, although this link remains unproven and warrants direct investigation. Taken together, these observations support a nuanced, context-dependent role for MLKL within the necroptosis pathway, warranting further investigation into how its metabolic effects can be therapeutically leveraged without unintended inflammatory or endocrine consequences.

Mechanistically, CRISPR/Cas9 deletion of MLKL in preadipocytes blunted induction of adipogenic genes, *Adipoq, Cepba* and *Pparg*, constraining lipid-loading during adipogenesis, in line with prior findings that MLKL deficiency impairs white-adipocyte differentiation partly via Wnt10b upregulation, a phenotype not reproduced by RIPK3 loss [23, 24]. Consistent with this, MLKL is required for white adipocyte differentiation but dispensable for beige adipocytes [24], aligning with our observation that MLKL loss limits lipid loading and promotes lipolysis in white fat. Notably, in differentiated adipocytes, siRNA-mediated MLKL knockdown produces smaller lipid droplets, lower neutral-lipid content, and greater glycerol release at baseline and during β-adrenergic stimulation, consistent with enhanced lipolysis and reduced lipid deposition. These cell autonomous changes parallel the *in vivo* reduction in adipocyte size and lower RER, implying that MLKL normally favours lipid storage whereas its suppression promotes lipid mobilisation and oxidation. Increased dark phase locomotor activity may further reinforce this lipid-first fuel shift. This lipolytic and oxidative bias fits within canonical pathways in which catecholamines activates adipose triglyceride lipase (ATGL) or hormone-sensitive lipase (HSL) via cAMP/PKA [26], while insulin restrains lipolysis through PI3K–AKT and phosphodiesterase-3B/α/β-hydrolase domain-containing 15 (PDE3B/ABHD15)-mediated suppression of Protein Kinase A(PKA) [52]. MLKL is positioned outside these classical networks, suggesting a role in lipid-droplet organisation, trafficking or mitochondrial interfaces – consistent with proposed non-canonical functions at membranes and organelles – to bias adipocytes toward storage rather than mobilisation [17-19, 41].

The preservation of insulin-stimulated AKT phosphorylation under TNF or palmitate stress indicates that MLKL knockdown buffers the insulin receptor-PI3K axis, potentially through reduced intrinsic lipid stress or altered membrane microdomains that coordinate insulin-signalling components [25]. Although increased lipolysis is expected to elevate circulating fatty acids and impair insulin action, whole-body glucose control improved. This aligns with reports that chronic β-adrenergic stimulation, which drives and enhances adipocyte oxidative capacity, can improve systemic glucose handling via coordinated adipokine signalling, substrate-selection adaptations, and inter-organ crosstalk. Improved glucose tolerance is also consistent with the recently described “adipoincretin” feedback loop, in which adipocyte PI3K/insulin signals to insulin secretion. Together, our data support a model in which cell-intrinsic protection of insulin–AKT signalling, combined with greater fatty-acid oxidation and smaller lipid droplets, lowers ectopic lipid and improves systemic insulin responsiveness.

## CONCLUSION

This study identifies MLKL as a regulator of adipocyte lipid handling whose sustained suppression restores metabolic flexibility and improves systemic glucose control independently of appetite reduction. By directly correcting adipocyte dysfunction that may persist despite incretin-based therapies, MLKL targeting represents a complementary strategy to address diet-induced obesity and insulin resistance. These findings expand the therapeutic landscape for metabolic disease and support further investigation of MLKL modulation – alone or alongside appetite-based interventions – within carefully defined tissue, dose, and safety parameters.

## CLINICAL PERSPECTIVES

- **Background:** Complications arising from obesity including insulin resistance are driven by adipocyte dysfunction and impaired metabolic flexibility, yet current therapies do not directly target adipocyte-intrinsic pathways regulating lipid handling and insulin signalling.
- **Summary of results:** Therapeutic MLKL suppression reduces adiposity, enhances lipid oxidation, and improves glucose homeostasis in obese mice by limiting adipocyte lipid storage, promoting lipolysis, and preserving insulin–AKT signalling under metabolic stress.
- **Significance:** These findings identify the RIPK1–MLKL axis as a clinically relevant therapeutic target to restore adipose metabolic health and insulin sensitivity, potentially complementary to appetite-based treatments for obesity and type 2 diabetes.

## Acknowledgement

We are grateful for the generous gift of antisense oligonucleotides targeting MLKL and the corresponding control provided by Dr Richard Lee and Dr Adam Mullick (Ionis Pharmaceuticals Inc., California, USA). We thank Prof Matthew Sweet (The University of Queensland) for helpful discussions on the manuscript. We also acknowledge Dr Sachini Forseka and Dr Alan Fang (The University of Queensland), Dr Vik Ven Eng (Monash University), and Drs Sam Young, Cathrine Hall, and Evelyn Leong (WEHI) for their valuable technical support.

## CRediT Author contribution

Conceptualization & Study Design: D.K. and J.M.M.; *In Vivo* Investigations and Data curation: X.L., Y.W., and D.K.; Tissue analysis, *In Vitro* Investigations and Data curation: M.S., H.X., A.A., S.F., M.A., K.D., S.N., T.R.; Supervision & Intellectual Input: M.S., B.G.D., J.M.M., Y-C.C., S.J.N. and D.K.; Writing – Original Draft: M.S. and D.K.; Writing – Review & Editing: M.S., S.J.N., B.G.D., J.M.M., and D.K.; Funding Acquisition: D.K.

## Data Availability Statement

The data that supports the findings of this study are available from the corresponding author upon reasonable request.

## FIGURE LEGENDS

**Supplemental Figure 1:**
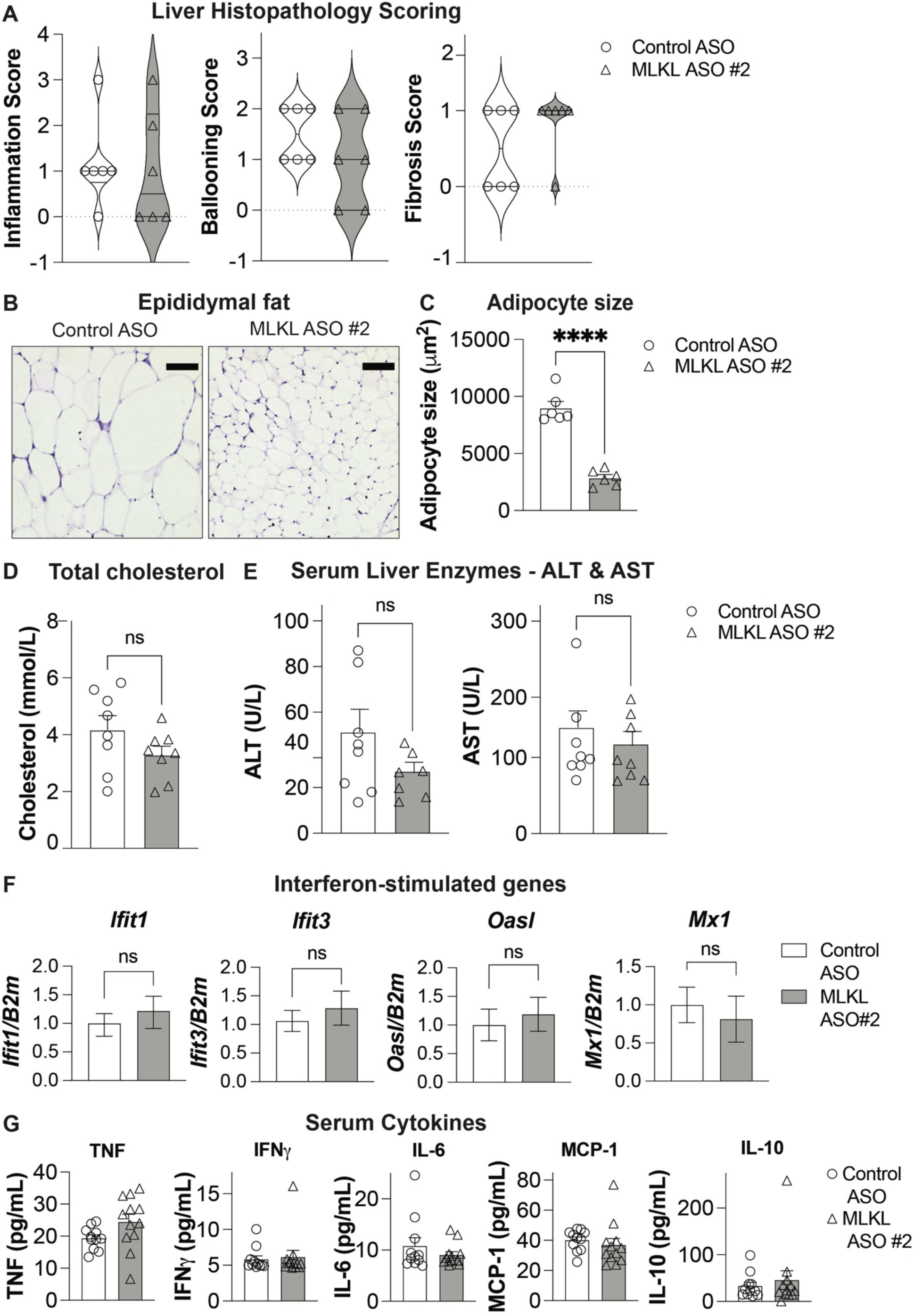
MLKL knockdown in obesity does not alter liver inflammation, fibrosis, hepatic injury markers or circulating cytokines despite reduced adipocyte size. Male C57BL/6J mice with diet-induced obesity were treated with control ASO or MLKL ASO#2 as described in Figure 1. **(A)** Liver histopathology scoring for inflammation, ballooning, and fibrosis. **(B)** Representative H&E-stained sections of epididymal adipose tissue. **(C)** Quantification of adipocyte size. **(D)** Fasted total serum cholesterol. **(B)** Serum alanine aminotransferase (ALT) and aspartate aminotransferase (AST). **(F)** Epididymal adipose tissue expression of interferon-stimulated genes (*Ifit1*, *Ifit3*, *Oasl*, *Mx1*), normalised to housekeeping gene, *B2m*. **(G)** Circulating serum cytokines (TNF, IFN-γ, IL-6, MCP-1, IL-10). Data are mean ± SEM. (n=12 mice/group). Significance assessed using a two-tailed Student’s *t*-test; ns, not significant; ****p<0.0001.

**Supplemental Figure 2:**
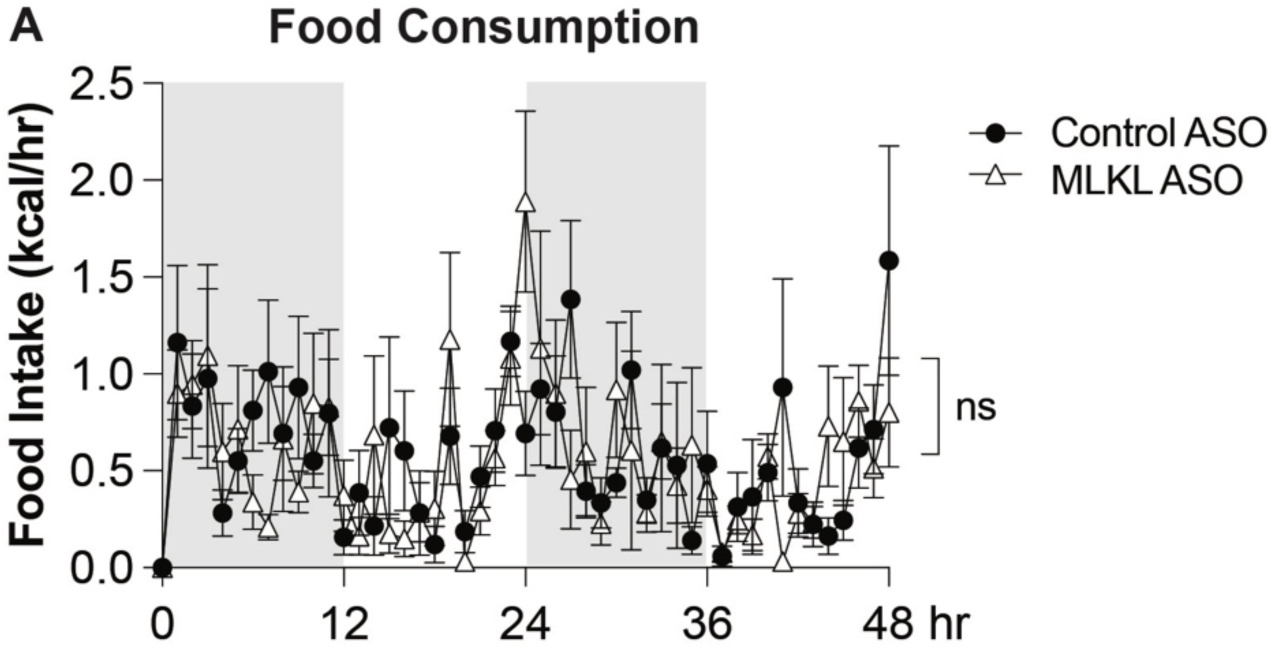
MLKL knockdown does not affect food consumption. Between weeks 16 and 17 of ASO-treatment, mice (n=8/group) were individually housed in the TSE Phenomaster indirect calorimetry system. (**A**) Food consumption over 48 h. Grey shading denotes the dark (night) phase and white shading denotes the light phase. Data were analysed using CalR beta software. Significance was assessed using two-way ANOVA with Bonferroni’s multiple comparison test, ns, not significant.

**Supplemental Figure 3:**
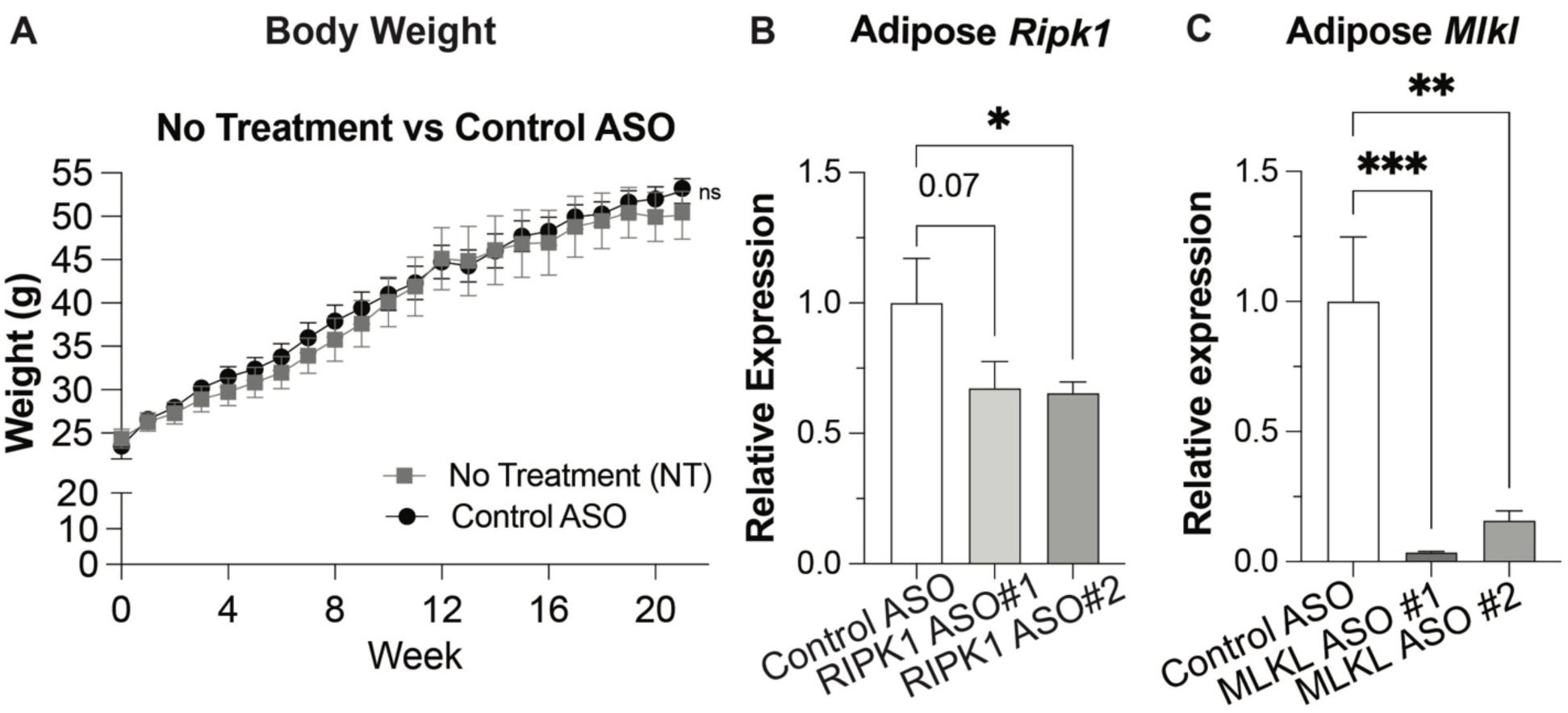
Control ASO does not affect body weight, while RIPK1 and MLKL ASOs effectively reduce adipose target gene expression. **(A)** Body weight trajectory comparing no-treatment (NT) and control ASO groups over 22 weeks. **(B)** Relative *Ripk1* mRNA expression in adipose tissue following treatment with control ASO or RIPK1 ASOs (#1 or #2). **(C)** Relative *Mlkl* mRNA expression in adipose tissue following treatment with control ASO or MLKL ASOs (#1 or #2). Data are mean ± SEM (n=8 mice/group). Statistical significance was assessed using one-way ANOVA with Dunnett’s multiple-comparison test; *p < 0.05, **p < 0.01, ***p < 0.001.

